# Synergistic short-term synaptic plasticity mechanisms for working memory

**DOI:** 10.1101/2025.05.07.652583

**Authors:** Florian Fiebig, Nikolaos Chrysanthisdis, Anders Lansner, Pawel Herman

## Abstract

Working memory (WM) is essential for almost every cognitive task. The neural and synaptic mechanisms supporting the rapid encoding and maintenance of memories in diverse tasks are the subject of an ongoing debate. The traditional view of WM as stationary persistent firing of selective neuronal populations has given room to newer ideas regarding mechanisms that support a more dynamic maintenance of multiple items. Various computational WM models based on different biologically plausible plasticity mechanisms have been proposed. We show that these proposed short-term plasticity mechanisms may not necessarily be competing explanations, but instead yield interesting interactions that broaden the functional range of models on a wide set of WM task motifs and simultaneously enhance the biological plausibility of spiking neural network models, in particular of the underlying synaptic plasticity.

While reductionist models (WM function explained by one particular mechanism) are theoretically appealing and have increased our understanding of specific mechanisms, they are narrow explanations. In this study we evaluate the interactions between three commonly proposed classes of plasticity, namely intrinsic excitability, synaptic facilitation/augmentation and Hebbian plasticity. We systematically test combinations of mechanisms in a spiking neural network model on a broad suite of tasks or functional motifs deemed principally important for WM operation, such as one-shot encoding, free and cued recall, multi-item delay maintenance and updating. Our analysis of the operational task performance indicates that a composite model is superior to more reductionist variants. Importantly, we attribute the observable differences to the principle nature of specific types of plasticity.

**Significance Statement:** Working memory - the ability to temporarily hold and manipulate information - is fundamental to cognition, yet the underlying neural and synaptic mechanisms remain debated. Traditional models propose and investigate isolated mechanisms like synaptic facilitation, often applied to specific tasks or rather simple delay maintenance. In contrast, this study proposes that broader working memory capabilities emerge from synergistic interactions among three commonly proposed short-term plasticity mechanisms: Hebbian learning, synaptic facilitation/augmentation and intrinsic excitability. Using biologically constrained spiking neural networks across a broad synthetic task battery, we show that combining mechanisms produces superior working memory capabilities compared with single isolated mechanism. This computational framework advances our understanding of how multiple plasticity mechanisms coordinate to support flexible, and robust working memory.

## Introduction

Working memory (WM) has attracted a lot of attention in brain science largely due to its central role in cognition. There is an ongoing multi-faceted dispute over the neural bases of WM, in particular about the involvement of so-called persistent activity in WM delay maintenance ^1,2^. This has rekindled interest in synaptic and neural plasticity mechanisms for short-term memory that can buffer information in neural and synaptic state changes without any neuronal spiking manifestations (activity silent) ^3–7^. It is tempting from a theoretical perspective to attribute working memory to one particular biophysically motivated short-term memory mechanism. History has seen many notable attempts ^8,9^, yet biology is not reductionist, and often relies on several mechanisms that act in concert to enable robust function across a range of task motifs^10^. Synaptic plasticity is diverse across different types of neurons and connections ^11^. Most of the observed and proposed short-term plasticity mechanisms of relevance to WM can be broadly grouped into three categories: presynaptic, postsynaptic and pre-post-synaptic.

Presynaptic plasticity-based WM is synaptic facilitation or augmentation ^12^, which transiently enhances synaptic efficacy via a calcium-mediated increase in synaptic release probability with dominating time constants up to 5s ^13,14^. Clear evidence for powerful synaptic augmentation in the medial prefrontal cortex was obtained using extracellular stimulation ^15^ and patch-clamp recordings ^13^. Consequently, there have been several models examining this synaptic WM hypothesis to explain WM delay maintenance ^16–21^ using a Tsodyks-Markram formalism ^22^, which is perhaps the most efficient and widely accepted phenomenological model of short-term plasticity (STP).

Postsynaptic mechanisms including rapid neuronal (i.e. intrinsic) excitability changes from one-shot learning, such as adaptation, plateau potentials, or graded increases in excitability ^23^ have been proposed as a cellular analog of WM ^24–26^ and address specific WM phenomena, such as serial order effects in computational models ^27–30^.

Pre-post-synaptic mechanisms are associative (i.e. Hebbian) and dependent on co-activity. This has long been held as the most important mechanism behind different forms of semantic and episodic long-term memory. Computational models often use variants of spike timing dependent plasticity (STDP) to implement Hebbian learning ^31^. Proposals that short-term Hebbian potentiation, such as NMDAR-STP may also be of importance to WM are less common ^32–34^, but increasingly part of the discussion ^35,36^. Theoretical neuroscience and recent reviews ^36,37^ have suggested fast Hebbian plasticity as a WM mechanism, even if it has been difficult to study in detail with current experimental technology, in part because LTP induction protocols and baseline monitoring may disrupt it ^38^. Neural network models capable of fast Hebbian learning generally support activity-silent WM with promising results in attractor memory networks, compatible with many electrophysiological and cognitive WM findings ^30,39–41^.

Synergies amongst plasticity mechanisms can enhance robustness, stability or functionality and performance in models. Combinations of Hebbian plasticity with secondary mechanisms can help stabilize runaway dynamics^42^, that would otherwise lead to implausibly high firing rates and excitatory postsynaptic potentials in spiking models, and may confer a degree of robustness and reliability to newly encoded memories ^43–45^. In addition, combinations of Hebbian plasticity with intrinsic excitability can simultaneously explain one-shot encoding and well-known serial order effects ^30,46^. Furthermore presynaptic facilitation/augmentation together with synaptic depression or neural adaptation synergistically gives rise to multi-item synaptic WM by making attractors quasi-stable (with short lifetime)^16,18,47,48^. Delay maintenance then relies on multiplexing a few memory items through related short-lived attractor reactivations in time, thereby periodically refreshing a small number of short-term memory traces. The same synergistic dynamic can be achieved with fast Hebbian learning instead of synaptic augmentation ^30,40^.

We aim to address two main issues we find with previous synergistic spiking models. Firstly, there are hardly any studies offering deeper insights into individual roles of these mechanistic components and how they substantiate emerging synergies. Secondly, computational models often gloss over the wide spectrum of WM functionality as described in consensus WM benchmarks^10^. Notable exceptions include work by Zenke et al. ^43^ investigating the ablation of individual mechanisms and Orhan et al. ^49^ investigating the multi-task capability of their plastic short-term memory model. Here we propose a more systematic approach to both issues, stressing the contributions of the three aforementioned categories (pre-, post-, pre-post-synaptic). We select a representative plasticity rule in each category and combine them in different constellations together with other biologically plausible activity-limiting mechanisms such as synaptic depression and neural adaptation in a spiking neural network model with cortical columnar architecture. In particular, we heavily exploit the Bayesian Confidence Propagation Neural Network (BCPNN) plasticity formalism as it inherently encapsulates associative Hebbian plasticity with intrinsic neuronal plasticity within an elegant phenomenological framework of Bayesian inference ^30,50–52^. The resulting WM models are thoroughly studied with biologically motivated constraints in mind, such as cell-type specific local connectivity, firing rates and the amplitudes of postsynaptic potentials. We compare functional capabilities and performance across a variety of WM task demands.

## Results

Our modeling approach relies on a spiking attractor memory neural network with a modular architecture and distributed representations, which has previously been used to model cortical memory ^30,53^ with a microcircuit model enabling various types of plasticity (see Methods, Fig.8, Fig.9). We use this model to simulate a range of WM tasks that provide insights into its various functional capabilities of key relevance for WM. By selecting one representative implementation for each of the 3 aforementioned categories of short-term plasticity mechanisms, we arrive at 7 meaningful plasticity configurations (Fig.1) to be examined across generic WM computational motifs (Fig.2), from one-shot novel item encoding, and multi-item delay maintenance with and without task disruption, to cued- and free (uncued, spontaneous) recall. The breadth of functional capabilities of WM models including the flexibility in dynamic adaptation to changing task demands, is often overlooked in computational studies primarily concerned with WM maintenance and recall, but may contribute explanatory power across different task scenarios. To address this gap we also investigate the updating of the task-relevant WM item set. The biophysical detail of the spiking model meanwhile enables us to constrain the model and assess biological plausibility on the basis of emergent electrophysiological dynamics, such as postsynaptic potentials and firing rates among others.

**Figure 1.**
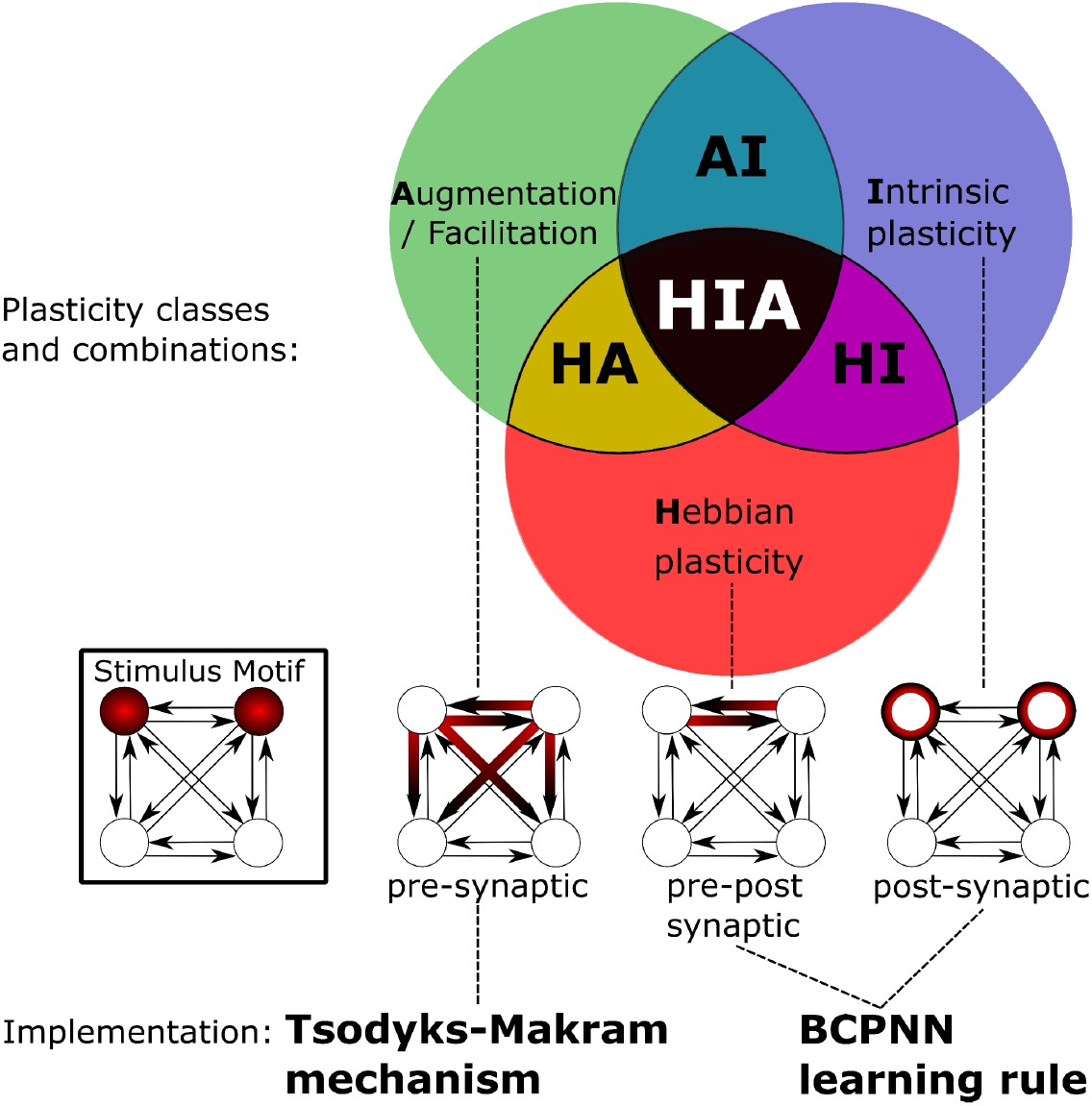
Plasticity Combinations. The Augmentation plasticity model is implemented using the well-known Tsodyks-Makram mechanism ^22^. The Bayesian Confidence Propagation Neural Network (BCPNN) learning rule implements intrinsic plasticity, as well as Hebbian plasticity. These 3 components can be simulated separately or together, yielding 7 scenarios to simulate and study (letters correspond to mechanism initials, e.g., **H** indicates Hebbian plasticity). Below, a simplified network schematic illustrates where different mechanisms may induce plastic changes at neurons and synapses in response to a spiking stimulus motif: **A**ugmentation changes with presynaptic activity, potentiating outgoing synapses of spiking neurons. **I**ntrinsic excitability changes with postsynaptic activity, increasing the excitability of spiking neurons. **H**ebbian learning at recurrent connections is driven by correlations in pre-post spiking.

**Figure 2.**
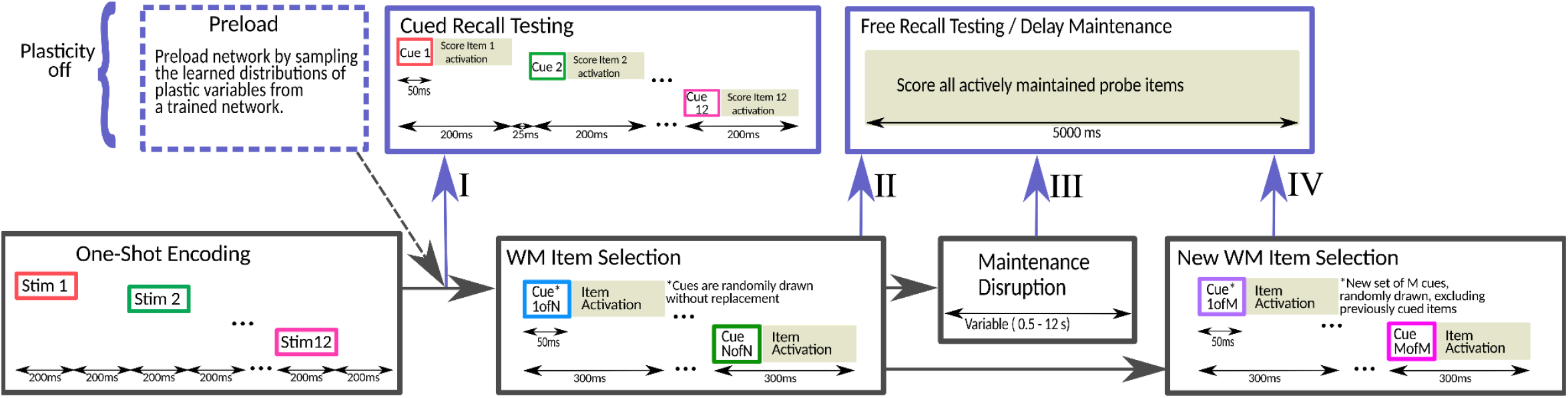
Overview of a general task structure simulated in different configuration scenarios. All simulated task configurations are sequential flows of memory operations (WM functional motifs) with ongoing plasticity (bottom row), and memory probes I-IV (top row), during which plasticity is typically disabled to assess the state of memory at that point without subsequent bias from rapid plastic changes induced by the testing protocol (*Methods - Stimulation Protocol, Memory Performance Tracking*). Stimulation is shown in item-colored boxes. Gray background denotes a free running network driven only by unspecific background noise. **One-shot Encoding** sequentially activates 12 neuronal assemblies to encode them as separate memory items into the network (acting as a reservoir of attractor memories). **Cued Recall Testing** activates all encoded items in turn by brief partial cues, checking for the stable recurrent activation of the full memory item thereafter. Probe I: probe all memory items. **WM Item Selection** uses cues to sequentially activate a subset of N unique items; cues are partial and brief to let the network activate the full item representation (entire neuronal assembly) on its own. Probe II: probe the selected WM items. **Maintenance Disruption** transiently lowers the background noise to disrupt Delay maintenance for a variable amount of time. Probe III: re-probe the WM items. **New WM item selection** repeats the previous item selection with M unique and novel items, drawn from the reservoir of 12 encoded items. Probe IV: probe new WM items.

**Figure 3.**
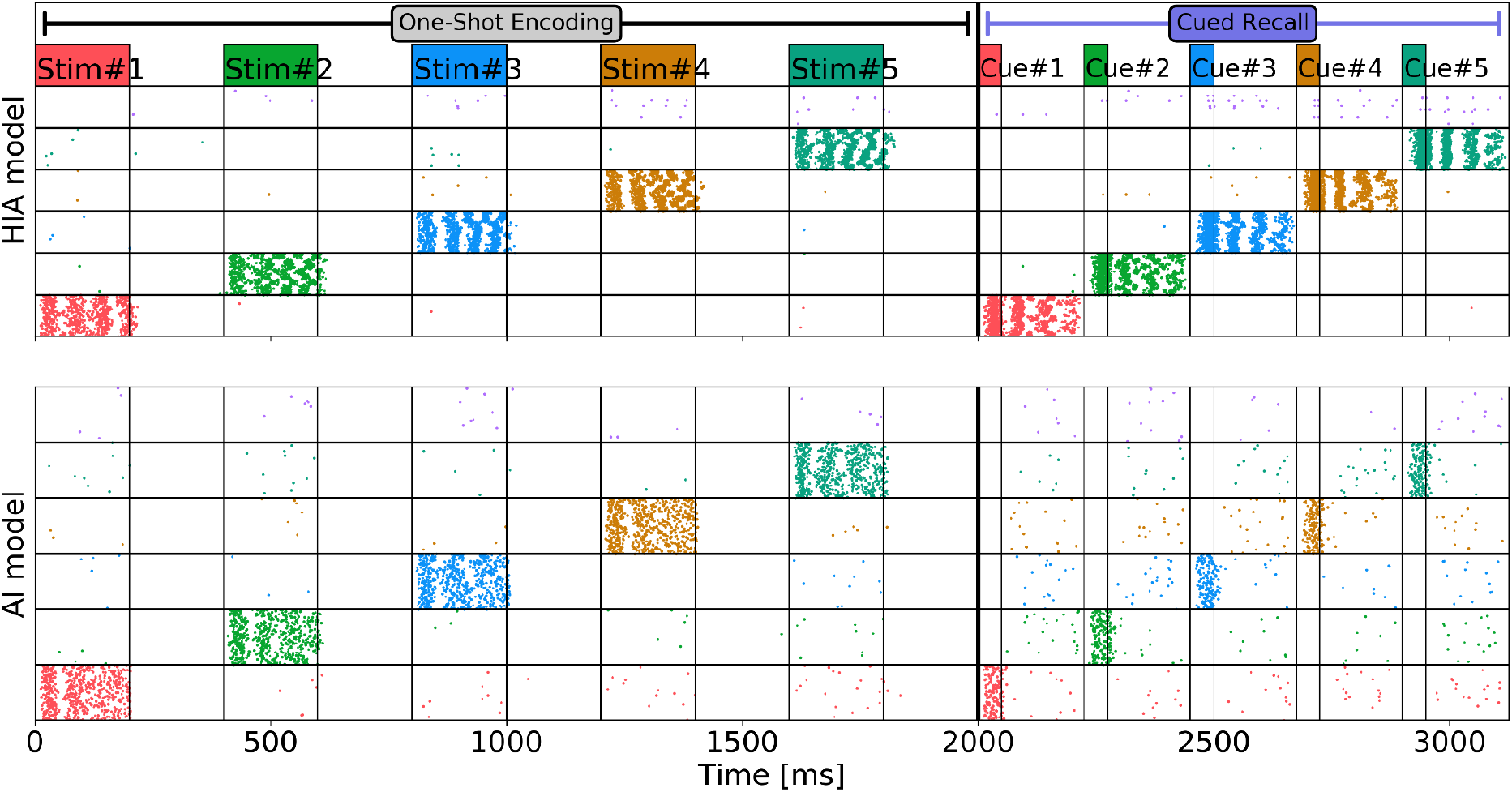
Subsampled, sorted spike rasters during One-Short Encoding and Cued recall testing epochs. Spikes are sorted and colored by their item-selectivity (only a subset corresponding to the same items during both simulated task epochs is shown). Stimuli and Cues are annotated in the same color on top. TOP: HIA model. BOTTOM: AI model (i.e. the same model, except without Hebbian plasticity).

### An overview of a general task structure and performance evaluation scenarios

To enable the study of WM operations, memory items have to first be encoded in the recurrent connections of the spiking network. Whilst many published models build on pre-encoded neuronal assemblies, we use the BCPNN learning rule to embed representations in an initially empty attractor memory network. So, prior to any WM operation, One-shot encoding (Fig.2) trains the model through the sequential stimulation of 12 non-overlapping stimuli (distributed sets of MCs encoding memory items, see Methods - Network Architecture and WM Neural Representations and Stimulation Protocol) for later retrieval or other WM operations. In order to save simulation resources and to enable some specific hypothesis testing later, we can also replace this fast online learning with a Preload, where all the network parameters affected by this initial learning are instead sampled from an already trained network, i.e. with encoded items (neural bias, synaptic AMPA and NMDA receptor conductances and the synaptic facilitation state; see Methods). Cued recall testing is used to examine the trained model. In particular, all the encoded items are probed in turn by brief partial cues (see Methods - Stimulation Protocol) in order to recall them (see Methods - Memory Performance Tracking and typical item activity in Fig.3). This way we evaluate a Cued Recall Score, the proportion of successfully recalled WM items.

To show how the network maintains several WM items at the same time, we select a small subset of memory items and track their activations over time. *WM item selection* (Fig.2) uses brief cues to sequentially activate (typically 4) memory items, similar to a set of sample cues in experimental WM tasks. We refer to this subset of activated items as the *WM set*. We then analyse the ongoing maintenance activity of the network during Delay maintenance / Free recall testing (Fig.2) to detect any memory item reactivations. Importantly, in this period, unlike during Cued recall testing, we do not provide any cues, but merely raise the unspecific background noise (see *Methods - Stimulation Protocol*). The observed memory item activations can be either interpreted as WM delay maintenance through WM replay ^54,55^ or as a so-called free WM recall ^46^, depending on the memory task paradigm. In both cases, we expect that only WM items belonging to the active WM set reactivate freely (uncued, spontaneously). To quantify this WM performance, we calculate a *Delay Maintenance Score*, the ratio of all successfully maintained (freely recalled) WM set items during Delay maintenance period over the total WM set size, corresponding to WM load (N in Fig.2). For further details on the identification of memory item activations and scoring, see *Methods - Memory Performance Tracking*.

We also test how robust the active maintenance of multi-item WM is by disrupting the delay period (Fig.2 - Maintenance disruption). We count all memory items as successfully retained after disruption if we then detect repeated free activations of the WM set items once we raise the background excitation again for a period of active Delay maintenance. The *Disruption Score* is the ratio of all successfully reactivated items over the total number of items in the current WM set (see *Methods - Memory Performance Tracking*).

To test the flexibility of the active maintenance we activate an entirely new WM set during *New WM item selection* (Fig.2). We expect the model to update its currently maintained memory content and switch to actively maintain new WM set items at the cost of the previous set items. This WM flexibility is evaluated by probing the active maintenance of the new WM set items with emphasis on the maintenance specificity wrt. the new updated WM set. The *Update Score* is the ratio of all successfully reactivated items from the new set over the total number of items in the new WM set (M in Fig.2). *Specificity* is the ratio of reactivations from the new WM set over the total number of reactivations during the delay period after the WM update (see Methods - Memory Performance Tracking), so unlike the *Update Score*, it accounts not only for the maintenance of the new WM set but also for how effectively the old WM set has been replaced relative the previous Delay maintenance.

### Cued recall reveals the need for Hebbian plasticity to encode novel items

Following One-shot encoding, we performed Cued recall testing in each plasticity model (cf. Fig.2). The full plasticity model featuring all three plasticity components (HIA) reliably retrieves all the encoded items. However, once the Hebbian plasticity component is removed (i.e resulting in the AI model) no WM items can be recalled. The spike raster in Fig.3. illustrates a simplified 5-item trial of encoding and subsequent Cued recall testing in both models. The stimulation drives the pyramidal cells in target minicolumns to fire in brief bursts. A similar pattern of activity reoccurs when we test for cued recall thereafter. Brief 50 ms cues to the previously stimulated populations quickly complete into the full spiking cell assembly, now driven by learned, recurrent connections rather than external stimulation. This activation persists until either neural adaptation and synaptic depression wears out the neuronal assembly, or a new stimulus or cue disrupts the ongoing reverberating activity through mutual inhibition (see Methods, Fig.9). Notably, the model without Hebbian plasticity shown in Fig.3 (bottom), does not achieve cued recall of any of the 5 items, even though the initial stimuli evoke a similar number of spikes during One-shot encoding.

In fact, the non-Hebbian models (I,A,AI) cannot perform cued recall following the prior attempt at One-shot encoding (see *Methods, Fig.10*). This is not a matter of tuning the models correctly, but simply because any recall of memory items as dynamic attractors in associative memory networks requires sufficiently strong associative connections to activate. This increase in excitation needs to be specific to the connections between neurons selective for the same item (Hebbian), rather than spread out (e.g. augmentation strengthens all presynaptic inputs). If the relevant synaptic connections are not preloaded (hard-coded) upon initialization of the network they instead need to be learned, yet non-Hebbian learning rules lack this ability ^36^. Previously published models of non-Hebbian synaptic WM typically rely on hard-coded associative connections establishing attractors for memory items prior to any WM test ^16,18^. If we are to compare non-Hebbian models with Hebbian models on such tasks, e.g. evaluating active delay maintenance, then we can first preload (frozen) associative connections from an otherwise identical network trained with Hebbian plasticity (see *Methods - Preload and Discussion - Preloading WM*).

### Compensating the unspecific excitability effects of synergy

The synergistic effect of different plasticity mechanisms naturally results in the enhanced excitability (or spiking activity) of the full HIA model, featuring all forms of plasticity examined in this work, when compared to reduced models. Removing entire plasticity mechanisms can shift the general excitability of the resulting models and in an attractor WM network model, this affects baseline recall performance even prior to any WM task. As we are not primarily interested in analyzing specific effects of plasticity on general excitability but rather in studying their implications for WM performance in different WM task paradigms, there is a need to establish a general baseline memory excitability across all the models so that a fair and insightful functional comparison can be made.

We have thus functionally compensated all incomplete models by replacing the missing activity-dependent plasticity components with fixed (non-plastic) initializations of the corresponding synaptic or neural parameters tuned to the minimum necessary level enabling cued recall of all the 12 baseline memory items. As explained in the previous section, non-Hebbian models (A,I,AI) are preloaded with fixed weights acquired through Hebbian learning in the same model with Hebbian plasticity. Furthermore, our implementation of intrinsic plasticity affects the excitability of neurons via a plastic neuronal current so the models without intrinsic plasticity (H,A,HA) are compensated via an extra constant neuronal current tuned to be sufficiently strong to enable cued recall (rather than setting this current to zero or some other unmotivated value). Finally, synaptic augmentation - as implemented by the Tsodyks-Makram mechanism - acts as a dynamic multiplier of synaptic efficacy. Accordingly, in all the model configurations without augmentation (H,I,HI) we replace this kind of plasticity with a fixed synaptic efficacy multiplier, adequate to enable cued recall. For further details see Methods - Preload. Having all the models compensated to perform cued recall of all memory items (Fig.2, Probe I), we can now focus on much more interesting comparisons in diverse WM task paradigms.

### WM item selection and delay maintenance (Probe II)

The ability to concurrently hold multiple task-specific WM items in an elevated active state is of general importance to WM function. We thus compare our models on their Delay maintenance performance (Fig.2, Probe II).

As we can see in the single trial example of Fig.4A, the HIA model can maintain all 4 initially selected task items through a short series of bursts over a delay interval of 5 s. The duration of these free WM item reactivations is limited by neural adaptation and synaptic depression. The multi-trial averaged *Delay Maintenance Score* at Probe II (Fig.4B) reveals that all models with synaptic augmentation (HIA,HA,AI,A) are able to reliably maintain most of the 4 selected items over the delay period, whilst the HI model often just maintains 2 of the 4 items, leading to a lower performance score. The H, and I models fail to maintain any WM set items (with the given model parameters), and disrupting Delay maintenance for a mere 2 s already incurs a dramatic loss of that limited maintenance performance (Fig.4C). We explore this in more detail in the section *Maintenance disruption*. Different models also show different maximum memory capacity during Delay maintenance, as we can see in Fig.4E. Overloading active maintenance by an excessive WM load may even decrease memory capacity.

**Figure 4.**
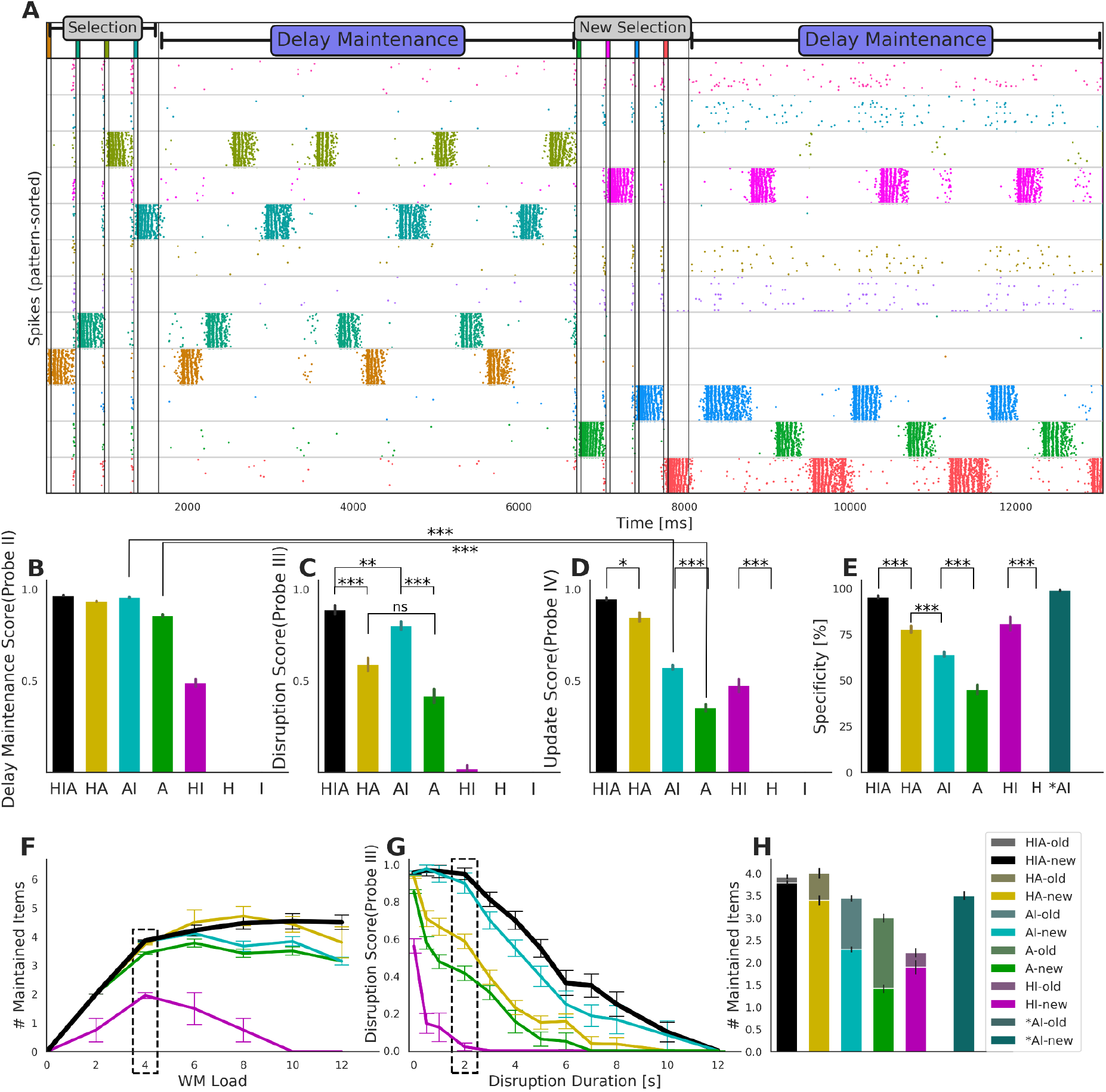
WM maintenance performance following WM item selection, disruption, and updating. **A:** Single trial of HIA model, sorted spike raster of WM item selection (4 items), subsequent 5 s Delay maintenance, New WM item selection (updating the WM set), followed by another period of delay maintenance. **B:** Delay Maintenance Score (Probe II) following WM item selection of 4 items for each of the 7 plasticity scenarios in the multi-trial average.** **C:** Disruption Score (Probe III) following WM item selection and 2 s of Maintenance disruption for each of the 7 models across trials.** **D:** Update Score (Probe IV) after New WM item selection for each of the 7 plasticity scenarios in the multi-trial average.** **E:** Maintenance Specificity after New WM item selection in the multi-trial average.** **F:** Load Saturation of the Delay maintenance. Datapoints mark the mean number of actively maintained items across trials at a given load in the multi-trial average.**The dashed marking annotates the 4-item-load default scored in B (relative to the load). **G:** Disruption Score over increasing duration of task disruption in all functioning models in the multi-trial average.**Disruption of 0 s denotes the control condition of immediate free recall (same data as **B**). The dashed marking annotates the 2 s disruption data (same data as **C**). **H:** Total number of items maintained during Delay maintenance after an update of the WM Set via New WM item selection, showing the multi-trial average**. 2-tone shading indicates items from the new WM set (colored according to the scenario), and items still maintained from the previous (old) WM set (grayed colors). *AI denotes an artificial hypothesis test based on the AI model (see next section) *\*\*All error bars indicate sem*.

### Updating and maintenance (Probe IV)

Beyond the general ability to maintain a limited number of rapidly selected task relevant memory items over some delay period, WM also needs to flexibly update its contents ^56,57^. We simulated this aspect of WM functionality with the New WM item selection (Fig.4D). The subsequent maintenance performance, quantified with the Update Score, reveals considerable differences between the models. While the HIA and HA models score high again, both the A (0.86 vs 0.35, p<1e-30, Mann-Whitney test) and AI models (0.96 vs 0.57, p<1e-48, Mann-Whitney test) drop their performance compared to the previous Delay Maintenance Score (cp. Fig.4D vs. Fig.4B). What do these performance differences tell us about the role of different plasticity mechanisms for flexible WM? It is helpful to break down the maintained items into 2 subgroups: those from the old WM set and those that should be maintained after the update with a new WM set (Fig.4H). We can see that the AI and A models (non-Hebbian) actively maintain 3 to 4 items, but often many of these maintained items actually belong to the old WM set. We measure this effect with the Specificity of maintenance (Fig.4E), the fraction of all reactivations during the delay period that belong to the new WM set. Apparently, maintenance may fail to fully shift to the new WM set, particularly in models without Hebbian plasticity. To show the network origins of this low Specificity of maintenance, particularly in non-Hebbian models, we examine the effects of plasticity after the update.

Since Hebbian plasticity is potentially strengthening the connections encoding individual items during New WM item selection, a first tangible hypothesis may be that items from the new set are more strongly encoded in the better performing Hebbian models (i.e. HA and HIA). Are new items simply more strongly encoded through increased recurrent excitation than old items in Hebbian models? To assess this, we can compute an effective “strength of the encoding” by averaging the contributions of all AMPA and NMDA synapses recurrently connecting pyramidal cells encoding the same item, filtering for items in the new WM set, and then performing the same analysis for items in the old WM set. It turns out, the ratio of relative recurrent excitation strength is generally very close to 1 in the multi-trial average (Fig.5-left), meaning new set items are not much stronger encoded than those in the old set. In particular, there is no obvious correlation between this strength ratio, and increased maintenance Specificity across trials and plasticity scenarios.

**Figure 5.**
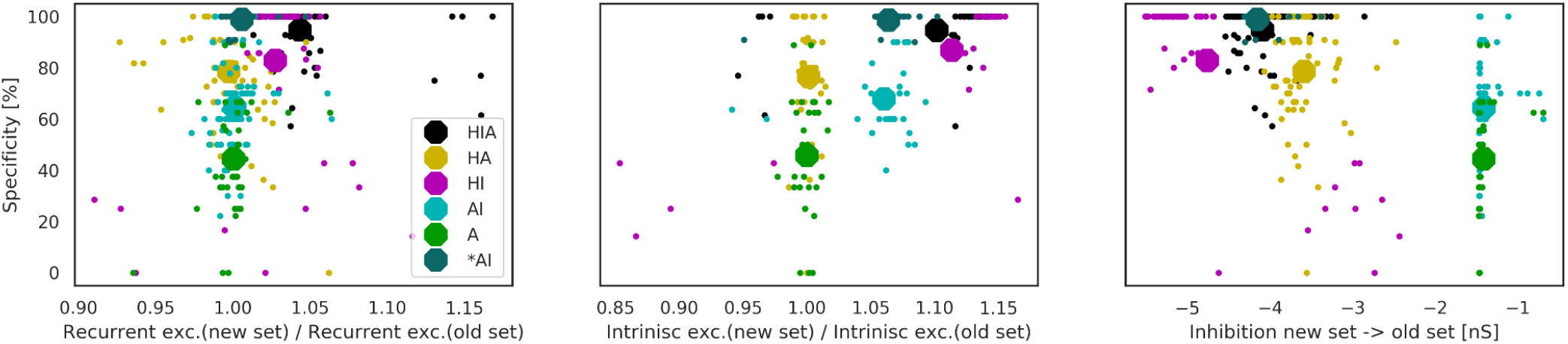
Trial-wise scatter plots of maintenance Specificity over three different metrics of learned network connectivity after WM updating. Each dot corresponds to an independent trial simulation. Its color marks the plasticity model used: HIA black, HA yellow, HI purple, AI cyan, A green, dark cyan denotes a special AI model with additional targeted inhibition. Larger octagons mark the multi-trial mean of trials with the same plasticity model. Left: Scatter over the ratio of recurrent excitation strength for cells selective for new and the old WM set items. Strength is based on the peak conductance transient between pyramidal cells sharing selectivity to the same item; the average is calculated over all recurrent connections of pyramidal cells selective for the same item and part of the same set). Middle: Scatter over the relative ratio of the average intrinsic excitability between items in the new WM set vs. items in the old WM set (averaged across all pyramidal cells selective for the same set). Right: Scatter over the average inhibition of cells selective for new set items on cells selective for old set items (measured as averaged across all pairwise connections).

Another aspect to consider is the beneficial role of intrinsic plasticity for the memory performance - models with intrinsic plasticity score higher (Fig.4D, compare H vs HI: 0.00 vs 0.47, p<1e-6; A vs AI: 0.35 vs 0.57, p<1e-10; and HA vs HIA: 0.85 vs 0.95, p=0.0194, Mann-Whitney test) and are more specific in their maintenance (Fig.4E, compare H vs HI: 0.00 vs 0.81, p<1e-8; A vs AI: 0.45 vs 0.64, p<1e-6; and HA vs HIA: 0.78 vs 0.95, p<1e-6, Mann-Whitney test). The mechanism can learn a cellular bias towards the most recently active items by boosting the intrinsic excitability of cells coding for new (i.e. recent) items. This is evident in the multi-trial average (Fig.5-middle), but still fails to explain why the HA model (which has fixed intrinsic excitability currents that are thus equal over the two sets) is significantly more specific in its maintenance than the AI model (0.78 vs. 0.64, p<1e-7, Mann-Whitney test). There is no obviously strong correlation between the stronger intrinsic excitability of new set items and higher Specificity during maintenance.

Yet one more possibility is that items in the new set actually suppress items from the old set in Hebbian scenarios, as the associative rule can learn disynaptic inhibition (see Methods). We can quantify this by averaging the learned connection strength from all neurons selective for items in the new set to all neurons selective for items in the old set (actually the learned peak conductances, see Methods). Indeed there is a large and significant difference between non-Hebbian and Hebbian models (average across models and samples: −1.41 vs −4.10, p<1e-49, Mann-Whitney test) and this targeted inhibition is negatively correlated with Specificity of the Delay maintenance (r=−0.60 (p<1e-30) across all models and simulations, see Fig.5-right). If we accept this as a hypothesis for why Hebbian models have higher Specificity, it still needs to be explained how this strong inhibition comes about and to demonstrate the causality of this effect on Specificity (and the Update Score).

The reason why the Hebbian model can learn such strong specific inhibition has to do with the temporal task structure. New WM item selection is a massed (rather than spaced) learning event, meaning the new items are activated in close temporal proximity (rapidly cued right after each other), leading to some amount of synaptic trace overlap (particularly with respect to the slow NMDA receptors), whilst spiking activity of the old WM set is temporally distant, and in fact anti-correlated from new item activity. The BCPNN learning rule learns excitation from spiking activity correlations and inhibition from anti-correlations (see *Methods - Spike-based BCPNN Learning Rule*). In our Hebbian models, this means that synaptic plasticity strengthens excitatory cell assemblies binding coactive cells (sharing the same item selectivity) and that neurons selective for items in the same set are also weakly exciting each o^ther. However BCPNN also learns inhibitory connections between neurons with anti-correlated activity, and the neurons with the most anti-correlated activity are selective for the other WM set items. One way we can see the spiking activity outcome of this fast Hebbian learning process is exemplified in Fig 3A after the updating the WM Set: Cells belonging to items in the old set have much lower background spiking activity than the cells selective for the new set items, or even the cells selective for the 4 completely unused (i.e. uncorrelated) items.

If this additional specific learned inhibition is causal (and sufficient) to the high *Specificity* in Hebbian models then synthetically adding only this targeted inhibition to an AI model should raise its *Specificity* to that of the complete model. To test this we identify the specific connections between cells selective for new set items and cells selective for old set items. We then sample the inhibitory weight values from such connections in the HIA model to set these particular connections in an AI model, without enabling Hebbian plasticity, and leaving everything else the same.

The result of this “transplantation” of structured inhibition dramatically boosts model *Specificity* to 97%, shown as the dark cyan *\*AI* model in Fig.4 and 5. The targeted inhibition effectively eliminates old set items from Delay maintenance (see Fig.4H). As new set items are not competing for reactivation with old items now, this results in an improved *Update Score* of 92%. This demonstrates the value of fast Hebbian plasticity in WM models, not only for rapid novel item encoding (Fig.3), but also for item set switching, as it occurs naturally in experimental WM tasks that include multi-item maintenance and repeated trials with different items.

### Maintenance disruption (Probe III)

The level of WM disruption has been shown to depend on the type of external interference ^58^. Due to the simplicity of our model, we confined our study to the case where interruption of the active maintenance of task relevant memory items in the delay period (replay of WM items) is operationalized as a transient reduction of background excitation to the spiking network to disable attractor activity (i.e. free, uncued recall; Fig.2 - Maintenance disruption) for some variable amount of time (.5 to 12 s). This enforces an activity-silent maintenance until background excitation is restored to normal levels.

We expect that a prolonged disruption should degrade the Delay maintenance performance of the model even after the end of the disruption, whereas short disruption should not have much of a lasting effect at all. This is because synaptic WM models generally do not require strictly persistent activity, and memory items are intermittently silent during Delay maintenance due to competition for foreground activity. A representative simulation trial can be seen in Fig.6. After activating a WM set of 4 items, we can see active maintenance of all items (*Delay Maintenance Score* of 1.0). Lowering the background excitation for 4 s disrupts this ongoing maintenance (see *Methods - Stimulation Protocol, Table 2*). When we raise the background excitation to its original value, only 2 memory items remain active during Delay maintenance leading to a *Disruption Score* of 0.5.

**Figure 6.**
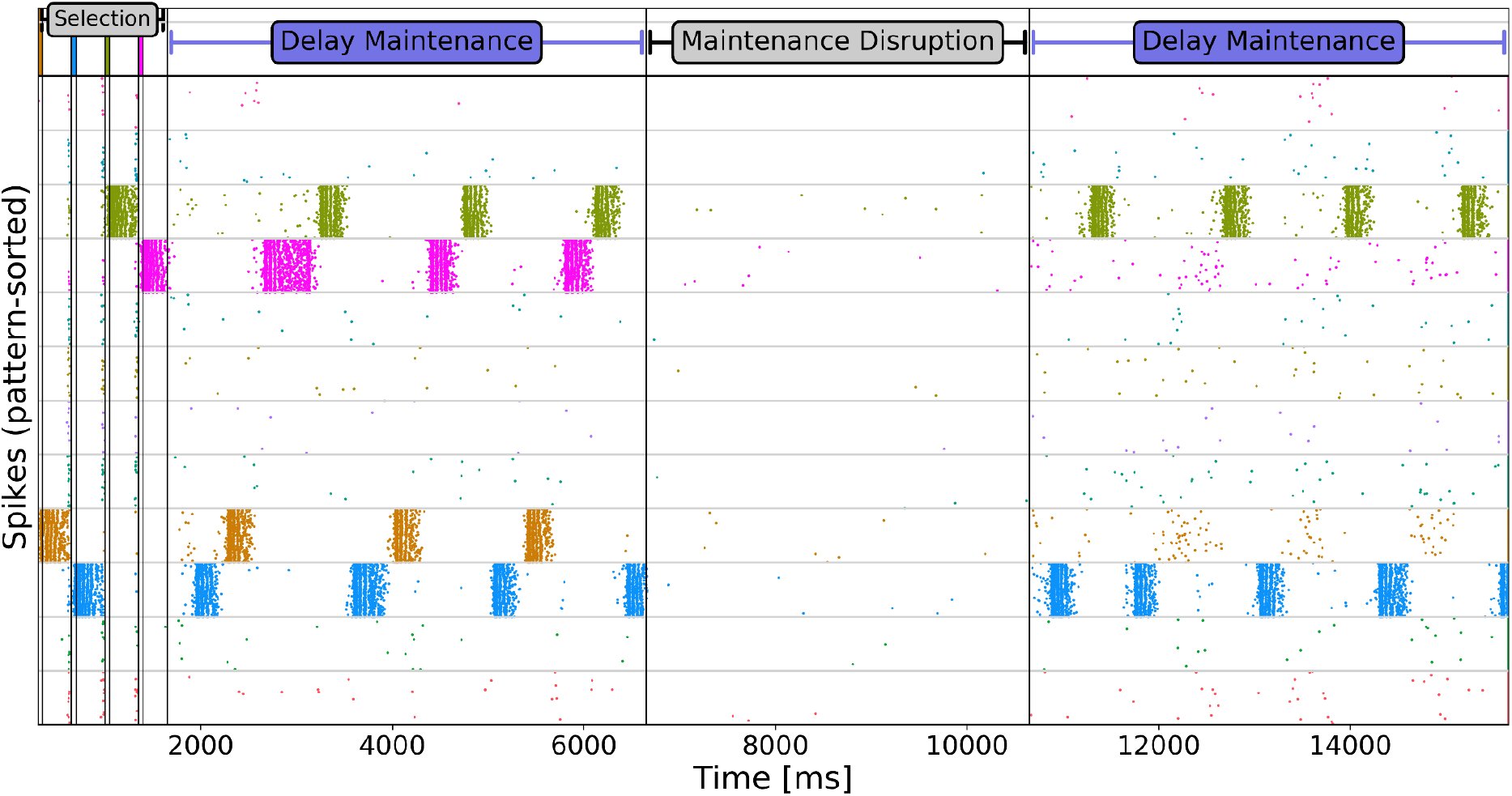
HIA model, Sorted spike raster of Delay maintenance with intermittent Maintenance disruption. Following the selection of 4 items and their successful maintenance, we disrupt the task by 4 s of intermittent silence in the network caused by considerably lower level of background excitation (uncorrelated noise). When we raise the background excitation back to the previous level, 2 of the 4 items are no longer being maintained in the subsequent delay period.

All plasticity mechanisms are based on neural and synaptic traces which generally decay with some time constant over the quiet period. The overall time constants for each mechanism is the same (by design), so we expect that the model can maintain WM for about that time constant, 5 s, but we can articulate some further expectations:

Firstly, free reactivations depend on many factors but are a somewhat thresholded process, so the most highly performing models (HIA and AI) can probably tolerate the longest periods of disruption and low-performing model, such as HI, may collapse after much shorter disruption. So overall, we do not expect the performance rank order of models to change much with disruption length.

Secondly, there is also an interesting aspect of Hebbian plasticity that may cause us to expect it to play a lesser role than other mechanisms here: Hebbian plasticity (as BCPNN) learns from spike correlations, which are not as strongly disrupted by silence, as they would be by uncorrelated noise. Neurons that are silent together, are in-fact correlated in their (in)activity, meaning correlation-based associative connection weights (H-type models) are probably not affected by silence as much as the more broadly activity-depended (A- and I-type models) that will react to the reduced pre- and post-synaptic activity, regardless of correlations. Because of the shared time-constant, we can expect a similar rate of performance loss from trace decay, in A- and I-type models then.

Fig.4G shows how much disruption each plasticity model can actually tolerate. The multi-trial averaged performance of each plasticity model degrades with the increasing duration of the disruption in the WM delay period. The combined model reliably preserves half of the task-relevant items even across 5 s long periods of delay Maintenance disruption. Confirming our expectations, the rank-order of models does not really change much. Models with higher Delay Maintenance Score can also tolerate the longest disruption. The presence of Hebbian plasticity does not strongly alter the Disruption Score (cp. A vs. HA: 0.62 vs. 0.69, p=0.07; AI vs. HIA: 0.76 vs 0.78, p=0.003). The effect of intrinsic plasticity is much more significant(cp. A vs. AI: 0.69 vs 0.76, p<1e-4; HA vs. HIA: 0.62 vs 0.78, p<1e-7). Two-way ANOVA reveals that the length of the disruption and the presence of intrinsic plasticity are both highly significant, while there is no interaction between the two.

### Matching Experimental EPSP Dynamics

We expect plausible WM models to be able to address a variety of WM tasks (as shown above). Because our model is primarily phenomenological we do not expect to make strong biophysical predictions, yet the model is detailed enough to invite attempts to link its electrophysiological properties to experimentally observable quantities, such as PSPs, that may help us identify further constraints on plausible plasticity models. Like other synaptic WM models, we predict that recently active items can be reactivated by a nonselective excitatory signal^59^.

The Bayesian Confidence Propagation Neural Network (BCPNN) learning apparatus ^30,41,50,51^ inherently encapsulates associative Hebbian plasticity with intrinsic neuronal plasticity within an elegant phenomenological framework of Bayesian inference. It learns excitatory and inhibitory postsynaptic potentials (EPSP/IPSP) between co-active and competing item-selective neurons. There is a stark difference between the performance of the HI and the HIA model across most task configurations however, such as Delay maintenance (Fig.4B). While the difference in baseline synaptic weights of those two models is small (see first spike in Fig.7-right, and also Methods - Fig.10), there is a big difference in the PSP dynamics during short spike bursts (often referred to as STP). The most common activity motif during maintenance is that of brief bursts of item-selective neurons with 3-7 spikes. Wang et al. ^13^ measured EPSP and their dynamics in prefrontal pyramidal cells with strong synaptic facilitation. We compare the dynamics of burst EPSPs (6 spikes at 20Hz) across different plasticity models with the biological record (Fig.7).

**Figure 7.**
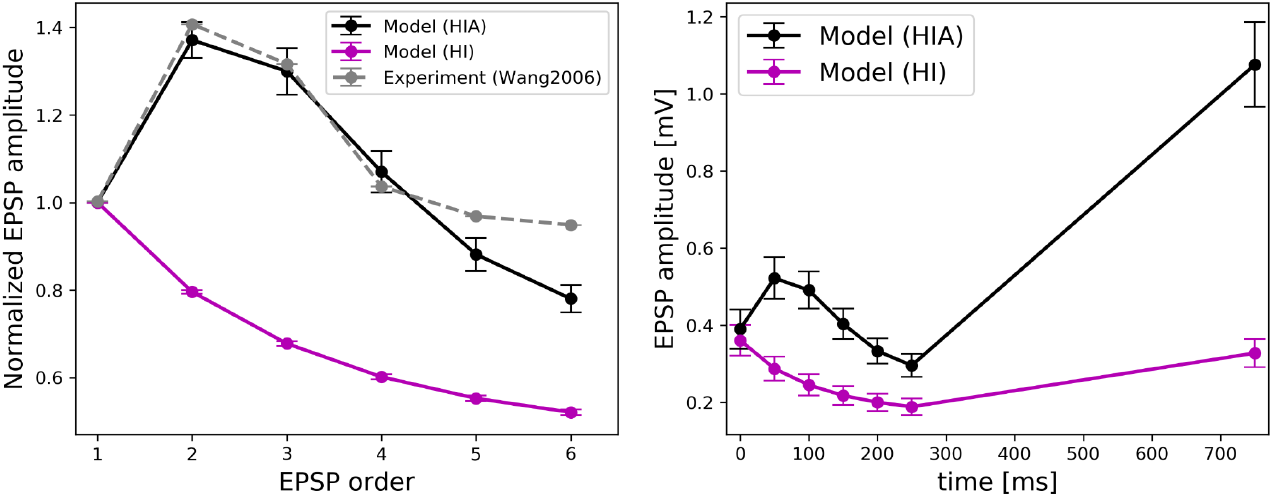
Excitatory Postsynaptic Potentials (EPSPs). Left: Normalized mean EPSP amplitudes in a 6-spike burst at 20Hz in two of the models and the relevant recordings by Wang et al. 2006. **Right:** EPSP mean amplitudes during the same stimulation protocol and subsequent and recovery test amplitude after 500ms. Error bars denote the 95% confidence interval.

As we can see in Fig.7 (left), the HIA model can reproduce the known short-term dynamics of facilitating prefrontal pyramidal cell EPSPs in ways the HI model cannot. The shape of the STP dynamic is influenced by synaptic facilitation and depression. The Tsodyks-Markram model implementing the decay of transmitter is governed by the ratio of remaining transmitter depleted with each transmitted spike (U=0.2 in our model, with Wang et al. reporting similar estimates), and the depression time constant (*τ*_rec_=280ms, as reported by Wang et al. (2006) for this class of synapses). Similar to the experimental record, augmenting synapses in the HIA model experience a roughly 40% increase in synaptic efficacy after the first spike, which then decays with the rapid depletion of transmitter across the burst (Fig.7 left). Because the synaptic depression decays away after some time, a recovery test spike 500 ms later will evoke a much larger EPSP if the time constant of facilitation/augmentation is longer than the depression time constant. Accordingly, the HI model without facilitation/augmentation recovers about 90% of its EPSP amplitude over 500ms, while the efficacy of HIA synapses increases to 176% above baseline amplitude (.4 mV to 1.1 mV). This is the reason why all our models with augmentation (A, HA, AI, HIA) perform much better at free recall (including Delay maintenance, see Fig.4B,C,D) than non-augmenting models (the average performance score for models with augmentation vs models without augmentation: 0.93 vs 0.31, p<1e-50, Mann-Whitney test). Long lasting augmentation (5s), after the end of a burst leaves attractors more “explosive” for a future item reactivation without raising the baseline EPSP. The latter would make HI models much more functional, as shown by a published HI-type model with 4-times larger baseline EPSPs closer to 1.5mV ^30^.

### Summary of key findings

In light of our results, **Hebbian plasticity** is a necessary component for novel encoding into an attractor WM model ^30,36^. Even when the novelty is just an association of long-term memory items, Hebbian plasticity is necessary to bind these, and has been used by Hebbian models to account for fast WM ^39,41^. Yet Hebbian plasticity alone is hardly sufficient. Under plausible biological constraints our own Hebbian-only (H) model achieves cued recall, but fails the majority of our Delay maintenance tasks in a straightforward manner from a general lack of excitability. Unpublished observations in the development of this model showed that the Hebbian only model can be restored to broader function by retuning the model, including specific violations of biological constraints, such as implausibly high background excitation, pyramidal firing rates larger than 30Hz, connection density (>20%) or strength (baseline pyramidal EPSPs>>1mV) outside the experimentally known distribution, stimulation durations exceeding typical WM tasks (larger than 1s), or more encoding repetitions (e.g. triple-shot encoding) enable active delay maintenance in the H model. A highly related HI-type model was previously shown to perform active delay maintenance with much longer stimuli which also resulted in larger EPSPs ^30^). We decided to forgo any such specific tuning of models in this study, because the fragility of reduced models is itself a valuable observation.

**Synaptic Augmentation** is key to effective, and biologically plausible active (bursty) maintenance in the attractor WM model. The Tsodyks-Makram implementation acts as a dynamic multiplier of the effective (learned) recurrent connectivity. This enables one-shot learning from brief stimuli, as it boosts the efficacy of synapses. Free reactivations enable active delay maintenance in our model. Without synaptic facilitation/augmentation, baseline EPSPs would need to be 3-4 times larger to achieve similar recovery spike amplitudes after short bursts (Fig.7, recovery spike EPSP), which are important to trigger attractor activations from noisy background activity. This is the main reason why models without synaptic augmentation do not perform well. Augmentation marries known EPSP constraints (i.e. sub 1mV baseline EPSPs ^13,60^) with attractor dynamics on a timescale relevant to WM, thus enabling a plausible attractor theory of WM in the first place. However, facilitation/augmentation by itself cannot encode novel items (Fig.3), is less robust against task disruption (Fig.4G), lacks capacity (Fig.4F), and is imprecise with the updating of actively maintained WM sets (Fig.4E,H).

**Intrinsic excitability** generally helps with stability, *Specificity* of maintenance (Fig.4E), and significantly improves performance across all our tasks (Fig.4B,C,D cp. A vs AI, H vs. HI, HA vs HIA), including robustness against longer task disruptions (Fig.4G). Our implementation of intrinsic excitability as part of the BCPNN learning rule was originally designed to match experimentally observed graded increases in excitability with repeated stimulation ^23,52^, but as with all other forms of plasticity there are many possible implementations of intrinsic excitability changes beyond simple spike-triggered adaptation that are outside the scope of this study, but could be be explored in future work.

## Discussion

We set out to explore the free combination of three different kinds of plasticity rules that have previously been proposed as WM mechanisms in a spiking neural network selected biological constraints, to evaluate whether there are specific synergies between them when applied to a broad battery of WM task requirements. After quickly finding that each additional mechanism contributes to the overall excitability of the network we normalized the general excitability such that all models would achieve cued recall of the encoded items (which also required the preloading of item encodings for non-Hebbian models). Even on that shared baseline we found important task-specific differences in the contributions of each of these plasticity mechanisms, which underpin the superiority of the fully combined model across the breadth of WM task motifs.

### Modeling choices

Our model follows the attractor theory of memory – a powerful paradigm for neural network WM models with reverberatory dynamics ^8^. Unlike popular bump attractor memory networks used to simulate visuospatial WM ^61^, our model operates in the space of discrete WM items, embodied by sets of cell assemblies wired by synaptic plasticity processes and activated upon cue-based WM recall or during short-term maintenance. This modelling framework helps explain, though not exclusively, a large body of experimental findings at the crossroads of neuropsychological and cognitive neuroscience domains. To reflect this, our generic model task flow accounts for functional operations needed to execute many real-world WM tasks in the spirit of goal-directed behaviour. The synergies we find in this regard are meaningful, because the combination of mechanisms allows us to address more than one task through a unified model, enabling an integrative rather than reductionist approach to addressing WM as a system with rich functionality. We do not account for the role of attention, executive control, short-term consolidation potentially supported by hippocampus or, from a broader perspective, the brainwide distributed nature of WM related phenomena ^62^. Instead, our focus is on core short-term memory network computations tentatively located in the prefrontal cortex, without explicit support of other WM-related circuits or subsystems. This model demonstrates that spiking models may be able to address the multi-task requirements of complex cognitive task batteries ^10^.

We acknowledge that there are many possible ways to classify and combine plasticity mechanisms. Our division of mechanisms by their activity factors (pre-, post-, pre-post-synaptic) conforms with well established nomenclature and focus on combinations of mechanisms that have previously been proposed as short-term memory mechanisms in their own right. We picked one representative from each class of short-term memory plasticity mechanisms to limit the scope, but one might choose others (i.e. STDP for Hebbian learning), or a more detailed calcium model for the presynaptic plasticity. We chose Tsodyks-Markram as our mechanisms for implementing the latter because it is the most commonly used model for synaptic short-term memory ^16^. For Hebbian plasticity, we opted for BCPNN over STDP because it already integrates Hebbian plasticity with intrinsic excitability in a principled way ^52^ and for its superior associative memory storage capacity ^51^. Any mechanism of the full model can be removed without causing any instability, enabling us to focus on the performance contributions of the learning rules to different WM tasks (see Results - Summary of key findings).

Alternative Hebbian learning rules (such as STDP) may require additional plasticity mechanisms to achieve stable weight and activity dynamics and avoid significant top-down interventions (like weight cropping, renormalization, network resetting etc.), and would likely involve some form of inhibitory learning corresponding to the di-synaptic inhibition learning inherent to BCPNN. BCPNN resolves this by being stable on its own. Whether for stability, robustness, biological plausibility or performance, the obvious solution is to combine Hebbian plasticity with other forms of fast plasticity ^43^. As our experiment with repeated WM item selection demonstrated, local Hebbian plasticity is also a candidate mechanism supporting some WM functions that are often hypothetically addressed at the systems level (see Rapid WM updating below). While Behavioral Time Scale Synaptic Plasticity (BTSP) is another associative learning rule that supports one-shot encoding and has significant experimental support, it is not strictly Hebbian, and has also been closely linked to hippocampal function, putting it outside the scope of this paper ^63,64^.

### Preloading WM

WM is multimodal, and leverages existing long-term memory ^65,66^ and sensory representations ^67^. Behavioural studies generally involve highly trained subjects/animals, which can be presumed to be highly familiar with the relevant stimuli and have stable representations. Many synaptic WM models ^47^ have used similar arguments to justify the preloading of the associative connectivity required for stable WM representations (cell assemblies/attractors). Hebbian WM models ^36^ provide an alternative to this preloading step by enabling the rapid formation of novel assemblies. We believe these views do not exclude each other. WM can predominantly operate on existing representations and also feature fast Hebbian plasticity, resulting in powerful models that may help address dynamic coding in WM. While we did not explore this aspect in depth here, previously published models have done so. By flexibly binding preexisting long-term memories, or selective populations through a dynamically recruited short-term memory (STM) population with flexible conjunctive cells ^39,68^, WM implements an indexing theory ^41^. Associated memories can be maintained or retrieved from the bidirectional connections of a rapidly learned index, which acts like a pointer, freeing up representational space for relevant task information, an aspect future models could explore in more detail. The memories themselves are presumed to be encoded in pre-existing, distributed representations, that may well span many brain areas and multiple modalities.

### Multiplexing WM in time

Multiplexing the maintenance of multiple items in time provides a natural explanation for the capacity limits of WM, which saturates at about 5 items in our model (see Fig.4F). There have been attempts to implement multi-item maintenance based on synaptic augmentation without multiplexing in time, but those results rely on dramatically implausible pyramidal firing rates to support the simultaneous persistent activation of multiple attractors in the same network ^17^. The idea of multiplexing multi-item maintenance in time is supported by macaque studies showing load-dependent increases in gamma burst rates during the delay period ^69^, and predicted by related models ^30^. Attractor dwell times emerge from the time-constants of synaptic depression and spike-frequency adaptation, and because there is a limited number of items that can be refreshed before memory traces decay too much, the WM overloading effects seen in our model (Fig.4F) are somewhat related to computational models based on the Time-Based Resource-Sharing theory ^70^.

### Rapid WM updating

Our study demonstrates the importance of rapidly learned (i.e. structured) di-synaptic inhibition for WM tasks that may require the rapid displacement of memory items through an active suppression of old content. Experimental evidence supports an integrated, simultaneous removal and encoding process, separate from passive forgetting that limits WM load and thus facilitates the maintenance of goal-relevant information by improving access to the remaining goal-relevant content ^56,57^. Our model captures this dynamic. While rapid updating is often addressed at the systems level as executive control (and we do not claim to address this here), our results indicate that some known Hebbian plasticity mechanisms (such as BCPNN) can support such memory operations without resorting to systems-level processes.

### Limitations

Our model is first and foremost a phenomenological model of memory dynamics and follows an attractor theory of memory with distributed neuronal assemblies for memory representations. Importantly, it lacks experimental verification down to the biophysical reality at the synaptic level. For a more realistic model, less constrained statistics of stimuli (overlapping or correlated random stimuli, spanning different coding levels, and complex protocols of presentation) should be considered. We did not address WM tasks that involve sequential information, but the issue was rigorously studied others ^71^. We also did not look into the rich oscillatory dynamics of this model, which have been strongly associated with WM function ^69^.

### Conclusion

Unlike what past reductionist models suggest, biology is likely to utilize many WM mechanisms in concert. Our study suggests that commonly used forms of plasticity proposed for the buffering of WM information besides persistent activity are eminently compatible, and yield synergies that improve function and biological plausibility. Combinations enable a more holistic model of WM responsive to broader task demands. Conversely, the targeted ablation of specific plasticity components reveals that different mechanisms are differentially important to specific aspects of WM function, advancing the search for more capable, robust and flexible models accounting for new experimental evidence of bursty and activity-silent multi-item maintenance. Importantly, we attribute the observable differences to the principle nature of specific types of plasticity. For example, we discover a previously undescribed synergistic function of Hebbian plasticity and synaptic facilitation/augmentation that enables the updating of multi-item WM sets through rapidly learned inhibition, which shifts active multi-item delay maintenance more effectively towards task-relevant representations than previously published spiking models. Our computational model simultaneously captures relevant electrophysiological, cognitive and behavioural effects, even though the synaptic plasticity rule is largely phenomenological. Further progress in understanding would benefit greatly from more quantitative experimental data on the dynamics of synaptic plasticity.

## Methods

We use the the NEST neural simulator ^77^ (see Code Accessibility), Model parameter values are found in Tab. 1-3.

### Neuron Model

We use an integrate-and-fire point neuron model with spike-frequency adaptation ^72^ which was modified by Tully et al. ^52^ for compatibility with a custom-made BCPNN synapse model NEST through the addition of the intrinsic excitability current. The model was simplified by excluding the subthreshold adaptation dynamics. Membrane potential *V*_*m*_ and adaptation current are described by the following equations:

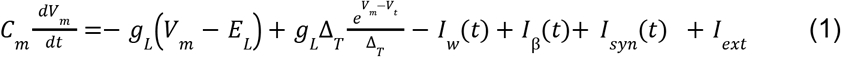

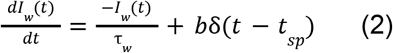

The membrane voltage changes through incoming currents over the membrane capacitance *C*_*m*_. A leak reversal potential *E*_*L*_ drives a leak current through the conductance *g*_*L*_, and an upstroke slope factor Δ_*T*_ determines the sharpness of the spike threshold *V*_*t*_. Spikes are followed by a reset of membrane potential to *V*_*r*_. Each spike increments the adaptation current *I*_*w*_ by *b*, which decays with time constant *τ* _*w*_. *I*_β_ is another intrinsic excitability current shaped by the Spike-based BCPNN Learning rule (see below). Simulated basket cells feature neither the intrinsic excitability currents *I*_β_ nor the the spike-triggered adaptation current *I* _*w*_.

Besides external input *I*_*ext*_ (*Stimulation Protocol*) neurons receive a number of different synaptic currents from their presynaptic neurons in the network (AMPA, NMDA and GABA), which are summed at the membrane accordingly:

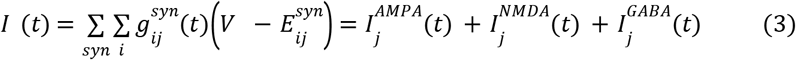

### Synapse Model and Augmentation Mechanism

Excitatory AMPA and NMDA synapses have a reversal potential *E*^*AMPA*^ = *E*^*NMDA*^, while inhibitory synapses drive the membrane potential toward *E* ^*GABA*^. Every presynaptic input spike (at 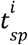 with transmission delay *t*_*ij*_) evokes a transient synaptic current through a change in synaptic conductance that follows an exponential decay with time constants *τ*^*syn*^ depending on the synapse type (*τ*^*AMPA*^≪ *τ* ^*NMDA*^).

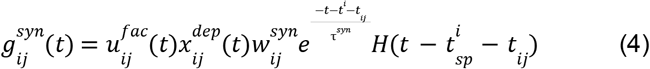

*H*(·) is the Heaviside step function. 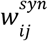 is the peak amplitude of the conductance transient, learned by the *Spike-based BCPNN Learning Rule* (next Section). Plastic synapses are also subject to STP according to the Tsodyks-Markram formalism ^73^, modeling the transmission-dependent synaptic facilitation/adaptation 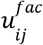 and the depletion of available synaptic resources 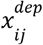 with a depression/reuptake time constant *τ*_*rec*_ and a facilitation time constant *τ* _*fac*_:

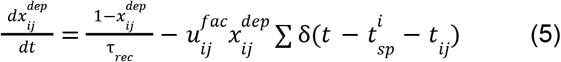

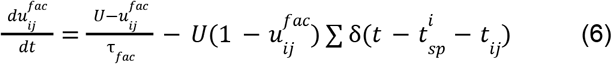

### Spike-based BCPNN Learning Rule (Hebbian and intrinsic plasticity)

Plastic AMPA and NMDA synapses are modeled to mimic NMDA-dependent Hebbian Short-term potentiation ^32,34^ with a spike-based version of the Bayesian Confidence Propagation Neural Network (BCPNN) learning rule ^52,74^. For a full derivation from Bayes rule, deeper biological motivation, and proof of concept, see Tully et al. (2014) and an earlier memory model implementation ^30^.

Briefly, the BCPNN learning rule makes use of biophysically plausible local traces to estimate normalized pre- and postsynaptic firing rates, as well as co-activation, which can be combined to implement Bayesian inference because connection strengths and neural unit activations have a statistical interpretation ^50,52^. Crucial parameters include the synaptic activation trace Z, which is computed from spike trains via pre- and postsynaptic time constants *τ*^*syn*^, which are the same here, but differ between AMPA and NMDA synapses: *τ*^*AMPA*^ = 5*ms, τ*^*NMDA*^ = 100*ms*

The larger NMDA time constant reflects the slower closing dynamics of NMDA-receptor gated channels. All excitatory connections are drawn as AMPA and NMDA pairs, such that they feature both components. Further filtering of the z-traces leads to rapidly expressing memory traces (referred to as p-traces) that estimate activation and co-activation:

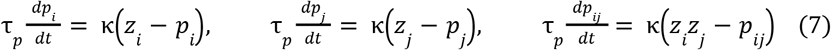

These traces maintain moving averages that support a palimpsest memory function. Short-term potentiation decay is known to take place on timescales that are highly variable and activity dependent ^75^.

The learning rule parameter κ (Eq.7), may reflect the action of endogenous neuromodulators, e.g. dopamine acting on D1 receptors that signal relevance and thus modulates learning efficacy. It can be used to switch off learning to fixate the network, or temporarily increase plasticity. In general, we trigger BCPNN plasticity concurrent with external stimulation, but switch it off during memory assessment, so that ongoing plasticity does not interfere with estimation of memory performance ^76^.

Tully et al. ^52^ showed that Bayesian inference can be recast and implemented in a network using the spike-based BCPNN learning rule. Prior activation levels are realized as an intrinsic excitability of each postsynaptic neuron, which constitutes a form of intrinsic plasticity, as these are derived from the postsynaptic firing rate estimate *p*_*j*_ and implemented in NEST as an individual neural current *I* _β_ with scaling constant β _*gain*_

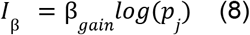

*I*_β_ is thus an activity-dependent intrinsic membrane current to the neurons, similar to the A-type potassium channel ^78^ or TRP channel ^79^. Intrinsic plasticity can be deactivated by inhibiting learning of *p*_*j*_ at the neuronal level, which freezes the intrinsic membrane current at its current value.

Synaptic weights are modeled as peak amplitudes of the conductance transient (Eq. 4) and determined from the logarithmic BCPNN weight, as derived from the p-traces with a synaptic scaling constant 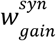.

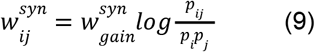

In this model, AMPA and NMDA synapses make use of 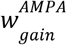 and 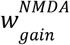 respectively. The logarithm in Equations 8,9 is motivated by the Bayesian underpinnings of the learning rule, and means that synaptic weights 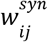 multiplex both the learning of excitatory and di-synaptic inhibitory interaction. The positive weight component is here interpreted as the conductance of a monosynaptic excitatory pyramidal to pyramidal synapse (Fig.9, plastic connection to the co-activated MC), while the negative component (Fig.9, plastic connection to the competing MC) is interpreted as di-synaptic via a dendritic targeting and vertically projecting inhibitory interneuron like a double bouquet and/or bipolar cell ^80–82^. Accordingly, BCPNN connections with a negative weight use a GABAergic reversal potential instead, as in previously published models of this kind ^30,52,83^. Model networks with negative synaptic weights have been shown to be functionally equivalent to those with both excitatory and inhibitory neurons with only positive weights ^84^. In the context of this particular model microcircuit and learning rule, this was explicitly and conclusively demonstrated by the addition of double bouquet cells ^85^. Code for the NEST implementation of the BCPNN synapse is openly available (see Code Accessibility).

### Network Architecture and WM Neural Representations

The model organizes cells into a grid of nested hypercolumns (HCs) and minicolumns (MCs), sometimes referred to as macrocolumns, and “functional columns” respectively. The Network is simulated with 16 HCs spread out on a rectangular grid. This spatially distributed network of columns has small conduction delays due to the distance between columns. Each of the non-overlapping HCs has a diameter of about 500 µm, comparable to estimates of cortical column size ^86^, contains 24 basket cells, and its pyramidal cell population has been divided into 12 MCs. This constitutes a sub-sampling from the roughly 100 MC per HC when mapping the model to biological cortex. We simulate 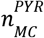 pyramidal neurons per MC to represent roughly the layer 2/3 population of an MC.

The distributed population code for each memory items is defined by a set of activated minicolumns across the networks hypercolumns (sparse due to WTA within each hypercolumn). This code is orthogonal, as we choose one active minicolumn in each hypercolumn to be selective for any given memory item without overlap between items (see Fig.8). The embedding of this code is achieved either via Hebbian learning during one shot item encoding or associative weights are preloaded at the start of the trial (see Fig.2, and Methods-Preload).

**Figure 8.**
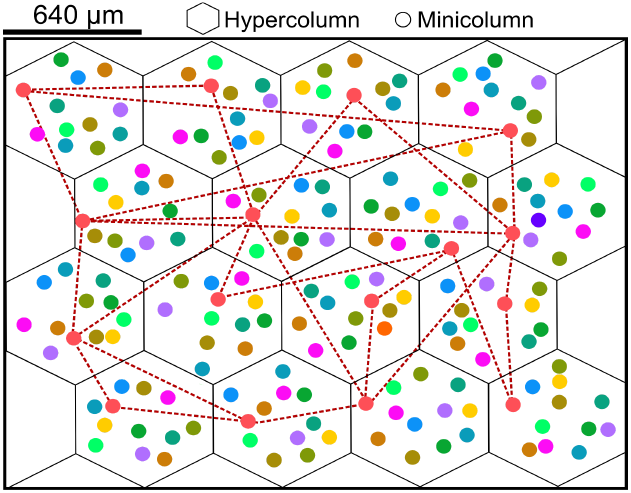
Basic modular network layout. The network is comprised of 16 hypercolumns (HCs), representing a small patch of neocortex. Each HC contains 12 minicolumns (MCs), which are composed of 30 pyramidal cells each, and preferentially active for 1 of the 12 orthogonal neuronal assemblies, as indicated by the 12 colors. For one neuronal assembly we also indicate the learning induced sparse excitatory long-range connections between subsampled pyramidal cells of similar selectivity

**Figure 9.**
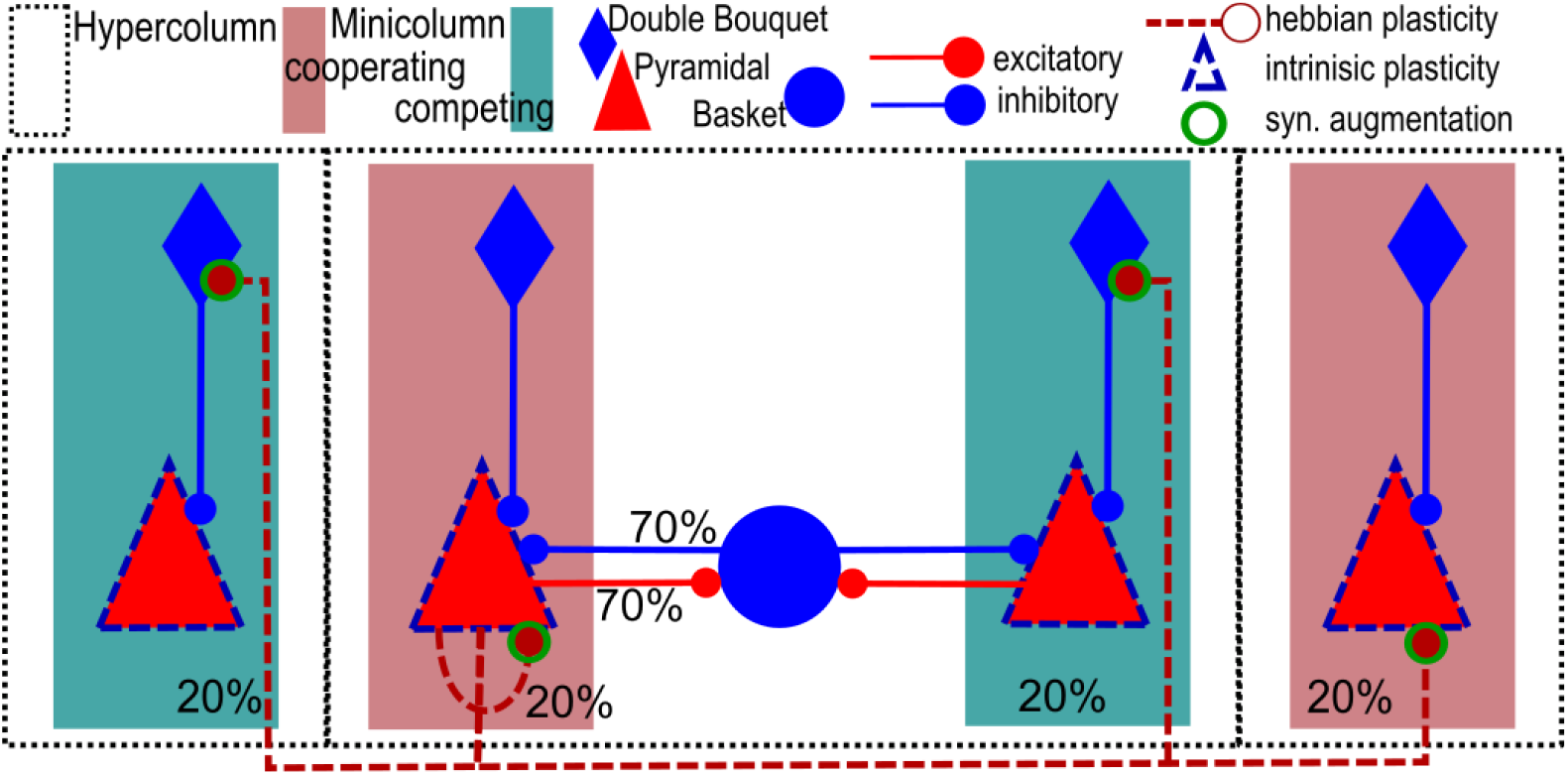
Columnar microcircuit and plasticity. Pyramidal cells connect recurrently within each MC, and share a pool of inhibitory basket cells within each HC, providing local feedback-inhibition to all pyramidal cells within the HC. Pyramidals also connect across HCs to other coactive or competing MCs. The strength of plastic connection (dashed in red) develops according to the Bayesian-Hebbian plasticity rule (BCPNN). Synaptic efficacy is modulated by facilitation/augmentation (dashed green), and the excitability of each pyramidal further influenced by the presence of intrinsic plasticity at each pyramidal cell (dashed blue). Connection probabilities are shown as percentages. Note that di-synaptic inhibition via DBC cells is not modeled directly, but instead expressed through the duplexing of excitatory and inhibitory learning in the BCPNN learning rule.

### Network Connectivity

Pyramidal cells project laterally to basket cells within their own HC via AMPA-mediated excitatory projections with a connection probability of *p*_*P*−*B*_, i.e. connections are randomly drawn without duplicates until the target fraction of all possible pre-post connections exist. In turn, they receive GABAergic feedback inhibition from basket cells (*p*_*B*−*P*_) that connect via static inhibitory synapses rather than plastic BCPNN synapses. This strong loop implements a competitive soft-WTA subnetwork within each HC ^87^. Local basket cells fire in rapid bursts, and induce alpha/beta oscillations in the absence of attractor activity and gamma, when attractors are present and active ^88^. Pyramidal cells form connections both within and across HCs at connection probability *p* _*pyr*_.

These projections are implemented with plastic synapses and contain both AMPA and NMDA components, as explained in subsection *Spike-based BCPNN Learning Rule*. Because BCPNN duplexes excitatory and inhibitory learning, inhibitory weights are expressed through a GABA receptor onto pyramidal cells, which indirectly models the existence of local double-bouquet cells, providing disynaptic inhibition onto pyramidal cells in their local columns. The existence of such local specificity of double-bouquet cell mediated inhibition was shown by DeFelipe et al. ^89^ and the equivalency of that indirect modeling approach confirmed by Chrysanthidis et al. ^85^.

In summary, the model thus features a total of 6.64 million plastic connections between its 5,760 simulated pyramidal cells, as well as 193,000 static AMPA- and GABA-mediated connections to and from 384 simulated basket cells and 192 double bouquet cells (virtual, one per MC), targeting all the pyramidal cells within their respective MC.

### Stimulation Protocol

The term *I*_*ext*_ in Equation 1 subsumes specific and unspecific external inputs. To simulate unspecific input from non-simulated columns, and other areas, pyramidal cells are continually stimulated with a zero mean noise background throughout the simulation. In each layer, 2 independent Poisson sources generate spikes at rate *r*_*bg*_, and connect onto all pyramidal neurons in that layer, via non-depressing conductances ± *g* _*bg*_. Before each simulation, we distribute the initial values of all plastic weights by a process of learning from 1000 ms of this unstructured background activity. A lower background rate (*r*_*bg*−*low*_) is used to disrupt active maintenance (Fig.4). During One-shot encoding we stimulate all 12 items in turn, such that Hebbian plasticity may encode the correlated firing structure of each neuronal assembly. To do this, we excite pyramidal cells belonging to component MCs of each distributed representation with an additional targeted excitatory Poisson spike train (rate *r*_*stim*_, length *t*_*stim*_, conductance *g*_*stim*_). This is followed by an inter stimulus period of the same length with lowered noise background rate *r*_*bg*−*low*_, followed by the stimulation of the next item, until all memory items have been stimulated once. To later cue the reactivation of any specific memory item (i.e. neuronal assembly/attractor) during Cued recall testing or WM item selection (Fig.4), we excite pyramidal cells belonging to a component MC with a brief excitatory Poisson spike train (rate *r*_*cue*_, length *t*_*cue*_, conductance *g*_*stim*_). We use partial cues defined by leaving out one of the coding minicolumns, which allows us to check for the associative completion of each activation as well. A brief 50 ms stimulus is usually sufficient to activate any previously encoded memory.

### Axonal Conduction Delays

We compute axonal delays *t*_*ij*_ between presynaptic neuron i and postsynaptic neuron j, based on a constant conduction velocity *V* and the Euclidean distance between respective columns. Conduction delays were randomly drawn from a normal distribution with mean according to the connection distance divided by conduction speed and with a relative standard deviation of 30% of the mean in order to account for individual arborization differences, and varying conduction speed as a result of axonal thickness/myelination. Further, we add a minimal conduction delay 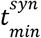 of 1.5 ms to reflect not directly modeled delays, such as diffusion of transmitter over the synaptic cleft, dendritic branching, thickness of the cortical sheet, and the spatial extent of columns:

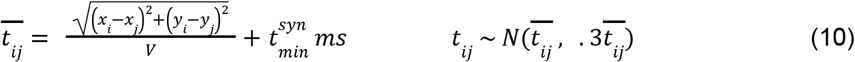

### Memory Performance Tracking

We track memory activity in time by analyzing the population firing rate of item-specific and network-wide spiking activity usually using an exponential moving average filter time-constant of 25 ms. Because the population code is orthogonal, and reactivations characterized by sizable gamma-like bursts, a simple threshold detector can extract candidate reactivation events and decode the activated memory. We further check for the complete activation of each assembly. Whenever targeted stimuli are used, we analyze peri-stimulus activation traces. When reactivation onsets are less predictable, such as during Delay maintenance, we extract reactivation candidates via a threshold detector trained at the 50^th^ percentile of the cumulative distribution of the population firing rate signal.

During Cued recall testing (Fig.4, Probe I), memory items are considered retrieved, if they fully activate within 175 ms following the cue for that item. The *Cued Recall Score* is then calculated as the ratio between the number of successfully retrieved items, and the total number of items cued (typically 12). During WM Item Selection and New WM Item Selection we cue the activation of a select WM set (typically 4 randomly selected items). New WM Item Selection excludes preciously selected patterns. During Delay maintenance (Fig.4, Probe II) there are no cues provided as items reactivate freely. Items are scored as successfully maintained if they activate at least twice. The *Delay Maintenance Score* is then calculated as the ratio between the number of successfully retrieved items, and the WM load (typically 4). The *Disruption Score* is computed the same way, except for the fact that the delay period only starts after a intermittent silence of variable length (.5 - 12 s). The *Update Score* is computed by tracking the activity of items in the updated WM set (typically 4) and then scored the same way. Notably, this excludes old WM items from the first set. Notably, we deactivate plasticity during the assessment of memory by setting *τ*_*fac*_ = 0 (Eq.6) and κ=0 (Eq.7). This is not because the model is unstable (BCPNN is stable, even with additional plasticity mechanisms), but because ongoing plasticity introduces a bias towards the memories, tested earlier, making it impossible to assess the state of the memory network without influencing subsequent task motifs. For example, if we do a cued recall test on any memory item with ongoing plasticity, we are more likely to see subsequent reactivations of that item if we then also test delay maintenance. By forgoing plasticity during memory assessment we can thus score multiple task motifs in a single continuous simulation run (Fig.4).

### Preload

Instead of training the network before each memory task, we may save time by preloading the network, which means sampling the values of plastic state variables from an already trained network. Each plasticity rules affects some of the network state variables (such as u, x, w, p-traces, etc.). As we discovered early on, incomplete models may fail to achieve basic cued recall (see Fig.3 and 10-D), because the inactivation of a mechanism freezes inappropriate initialization values (Fig.10 A-C), lowering overall excitability. Rather than re-tuning background excitation for each model, preload offers a means to compensate for this by loading appropriate values. In order to capture the variability of the learned distribution of such network state variables in dynamic spiking neural networks, we can even preload state variables from learned and stored distributions of these parameters. If a model does not have the specific mechanism, the affected variables will obviously be frozen after the preload.

Intrinsic plasticity for example affects neuronal excitability through a change in a neuron-specific membrane current *I*_β_ (Eq.1), which is itself dependent on the learned neuronal p-trace (Eq.8). If we disable intrinsic plasticity in any model, this means freezing the strength of this current, such that it no longer changes with activity but what value to freeze the underlying p-trace at, is not self-evident. So in the example of the HA model without intrinsic plasticity we preload the underlying distribution of neural p-traces from the full HIA model, which then determine a distribution of bias currents *I*_β_ (Eq.8). We then set a bias gain β _*gain*_ = 35*pA*, that enables cued recall of all items. Similarly, non-Hebbian models are preloaded with frozen attractor weights (no synaptic gain chain is necessary to achieve cued recall here), and models without augmentation have their synaptic gains (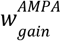 and 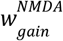) boosted by 50%, which allows all non-augmentation models to achieve cued recall. This compensates the reduced models for the loss of overall excitability (Fig.10-E), but not more than necessary for cued recall of all encoded items and thus enables a more interesting comparison of the 7 different plasticity combinations (Fig.1) across the more dynamic WM tasks thereafter (Fig.2).

**Figure 10.**
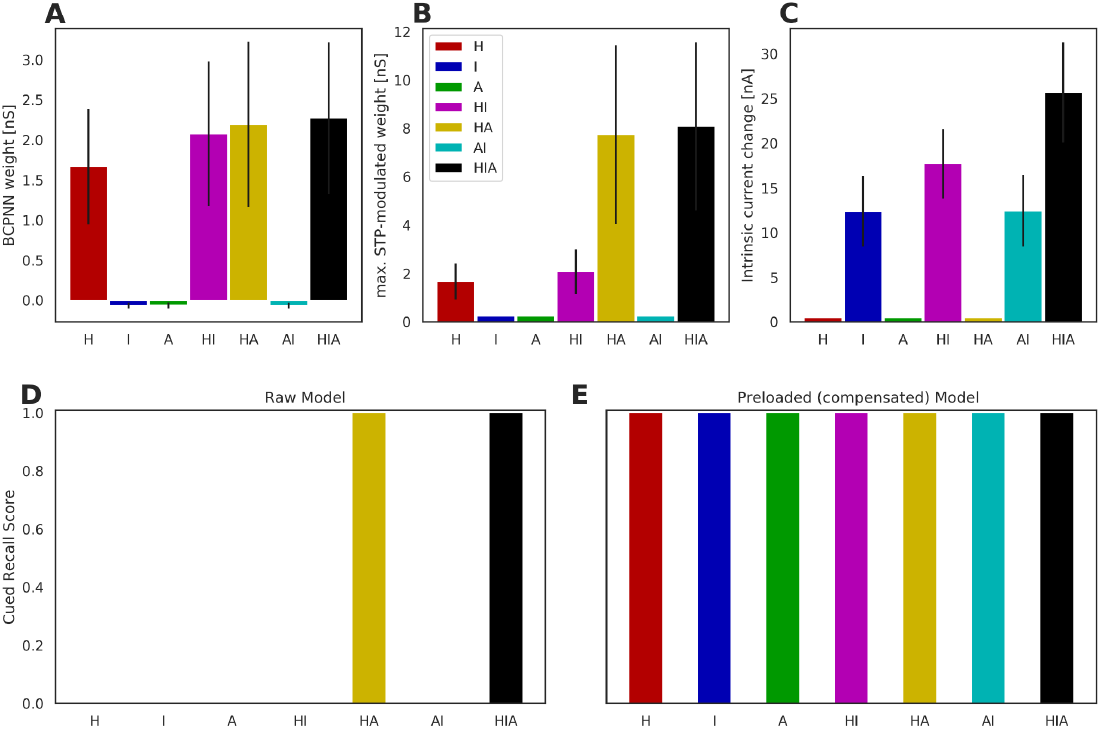
**A:** BCPNN weight of excitatory recurrent attractor connections (combined across AMPA and NMDA receptors) following 1-shot-encoding, averaged over all excitatory item assembly connections and trials. Error bars denote standard deviation over all excitatory item assembly connections and trials. **B:** STD-modulated maximal weight (max. amplitude of the conductance transient of excitatory recurrent attractor connections) including the multiplicative effect of augmentation) following 1-shot encoding of excitatory recurrent attractor connections, averaged over all excitatory item assembly connections and trials. Error bars denote standard deviation over the same. **C:** Absolute increase in intrinsic current (nA) of item-selective neurons following 1-shot encoding, averaged over all neurons in item assemblies and trials. Error bars denote standard deviation over all neurons in item assemblies and trials. **D:** Cued Recall Score (Probe I) prior to compensation through preload. No variability across at least 20 trials per scenario. **E:** Cued Recall Score (Probe I) after compensation through preload, constituting the baseline for all further model comparisons, averaged over at least 20 trials per scenario. No variability in *Cued Recall Score* detected, as all trials succeed in retrieving all.tested items.

## Acknowledgements

This research was supported by Vetenskapsrådet 2018-05360 and 2018-07079, the Swedish e-Science Research Centre (SeRC), Digital Futures, and European Commission, Directorate-General for Communications Networks, Content and Technology (grant no. 101135809). The simulations were performed on resources provided by National Academic Infrastructure for Supercomputing in Sweden (NAISS) at the PDC Center for High Performance Computing, KTH Royal Institute of Technology.

## Code Accessibility

We use the NEST simulator^77^ version 2.2 for our simulations, running on a Cray XC-40 Supercomputer of the PDC Centre for High Performance Computing. The custom-build spiking neural network implementation of the BCPNN learning rule for MPI-parallelized NEST is freely available on github: https://github.com/Florian-Fiebig/BCPNN-for-NEST222-MPI and the microcircuit model also available on ModelDB: modeldb.yale.edu/257610. Model output data and python scripts for generating all related figures from it are available on request.

## Parameter Tables

**Table 1.**
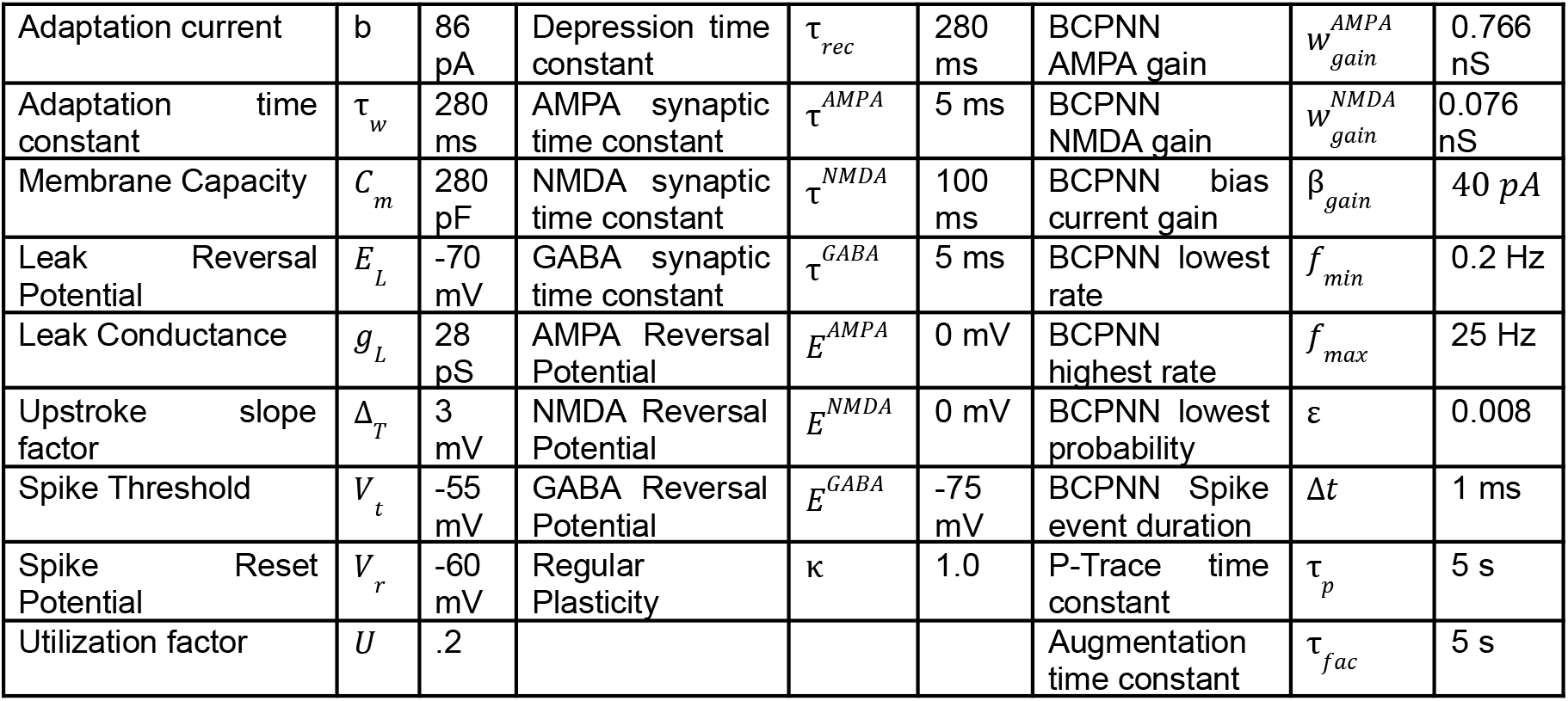
Neurons, synapses, and plasticity.

|  |  |  |  |  |  |  |  |  |
| --- | --- | --- | --- | --- | --- | --- | --- | --- |
| Adaptation current | $b$ | 86 pA | Depression time constant | $\tau_{rec}$ | 280 ms | BCPNN AMPA gain | $w_{gain}^{AMPA}$ | 0.766 nS |
| Adaptation time constant | $\tau_w$ | 280 ms | AMPA synaptic time constant | $\tau^{AMPA}$ | 5 ms | BCPNN NMDA gain | $w_{gain}^{NMDA}$ | 0.076 nS |
| Membrane Capacity | $C_m$ | 280 pF | NMDA synaptic time constant | $\tau^{NMDA}$ | 100 ms | BCPNN bias current gain | $\beta_{gain}$ | 40 pA |
| Leak Reversal Potential | $E_L$ | -70 mV | GABA synaptic time constant | $\tau^{GABA}$ | 5 ms | BCPNN lowest rate | $f_{min}$ | 0.2 Hz |
| Leak Conductance | $g_L$ | 28 pS | AMPA Reversal Potential | $E^{AMPA}$ | 0 mV | BCPNN highest rate | $f_{max}$ | 25 Hz |
| Upstroke slope factor | $\Delta_T$ | 3 mV | NMDA Reversal Potential | $E^{NMDA}$ | 0 mV | BCPNN lowest probability | $\varepsilon$ | 0.008 |
| Spike Threshold | $V_t$ | -55 mV | GABA Reversal Potential | $E^{GABA}$ | -75 mV | BCPNN Spike event duration | $\Delta t$ | 1 ms |
| Spike Reset Potential | $V_r$ | -60 mV | Regular Plasticity | $\kappa$ | 1.0 | P-Trace time constant | $\tau_p$ | 5 s |
| Utilization factor | $U$ | .2 | | | | Augmentation time constant | $\tau_{fac}$ | 5 s |

**Table 2.** Network size, Conduction delay, Stimulation.

|  |  |  |  |  |  |
| --- | --- | --- | --- | --- | --- |
| STM patch size | 2 x 2 mm | | Background activity rate | $r_{bg}$ | 1190Hz |
| Simulated HCs | $n_{HC}$ | 16 | Low Background activity rate | $r_{bg-low}$ | 870Hz |
| Simulated MC per HC | $n_{MC}$ | 12 | Background conductance | $g_{bg}$ | $\pm 1.5$ nS |
| Hypercolumn diameter | $d_{HC}$ | 0.5 mm | Targeted Stimulation rate | $r_{stim}$ | 350 Hz |
| Layer 23 pyramidal per MC | $n_{MC}^{PYR}$ | 30 | Targeted Stimulus conductance | $g_{stim}$ | +3.0 nS |
| Basket cells per MC | $n_{MC}^{basket}$ | 2 | Targeted Stimulus duration | $t_{stim}$ | 200 ms |
| Axonal Conduction Speed | $V$ | .2 $\frac{m}{s}$ | Cue stimulus duration | $t_{cue}$ | 50 ms |
| Minimal conduction delay | $t_{min}^{syn}$ | 1.5 ms | Cue stimulation rate | $r_{cue}$ | 375 Hz |

**Table 3.** Projections.

| Source | Target | Type | Symbol | Value |
| --- | --- | --- | --- | --- |
| Pyramidal | Basket | probability | $p_{P-B}$ | 0.7 |
| Pyramidal | Basket | conductance (static) | $g_{P-B}$ | +3. nS |
| Basket | Pyramidal | probability | $p_{B-P}$ | 0.7 |
| Basket | Pyramidal | conductance (static) | $g_{B-P}$ | -7 nS |
| Pyramidal | Pyramidal | probability | $p_{pyr}$ | 0.2 |
| Pyramidal | Pyramidal | AMPA gain (BCPNN) | $w_{gain}^{AMPA}$ | 0.766nS |
| Pyramidal | Pyramidal | NMDA gain (BCPNN) | $w_{gain}^{NMDA}$ | 0.076nS |

## Notes

### Competing Interest Statement

The authors have declared no competing interest.

### Summary of Updates

Updated for more concise language, and some more recent, relevant references, particularly in the Introduction

